# Large-scale comparative genomics unravels great genomic diversity across the *Rickettsia* and *Ca*. Megaira genera and identifies Torix group as an evolutionarily distinct clade

**DOI:** 10.1101/2021.10.06.463315

**Authors:** Helen R Davison, Jack Pilgrim, Nicky Wybouw, Joseph Parker, Stacy Pirro, Simon Hunter-Barnett, Paul M Campbell, Frances Blow, Alistair C Darby, Gregory D D Hurst, Stefanos Siozios

## Abstract

*Rickettsia* are intracellular bacteria originally described as arthropod borne pathogens that are emerging as a diverse group of often biologically important, non-pathogenic symbionts of invertebrates and microeukaryotes. However, sparse genomic resources for symbiotic strains and for the sister genus (*Candidatus* Megaira) inhibit our understanding of *Rickettsia* evolution and biology. Here, we present the first closed genomes of *Ca*. Megaira from an alga (*Mesostigma viride*), and Torix *Rickettsia* from midge (*Culicoides impunctatus*) and bed bug (*Cimex lectularius*) hosts. Additionally, we sequenced and constructed draft genomes for *Ca*. Megaira from another alga (*Carteria cerasiformis*), Transitional group *Rickettsia* from tsetse fly (*Glossina morsitans submorsitans*), and Torix *Rickettsia* from a spider mite (*Bryobia graminum*). We further extract 22 draft genomes from arthropod genome sequencing projects, including 1 Adalia, 4 Transitional, 1 Spotted Fever, 7 Torix, 7 Belli and the first Rhyzobius and Meloidae *Rickettsia* group genomes. We used new and existing *Rickettsia* genomes to estimate the phylogeny and metabolic potential across groups and reveal transitions in genomic properties. These data reveal Torix as unique amongst currently described *Rickettsia*, with highly distinct and diverse accessory genomes. We confirm the presence of a third subclade of Torix, previously only known from gene marker sequences. Further, Torix share an intact pentose phosphate pathway with *Ca*. Megaira, not observed in other *Rickettsia*. Considering the distinctness and diversity of Torix, we propose that the group be named *Candidatus* Tisiphia. The wide host range of *Ca*. Tisiphia symbionts necessitates onward research to understand the biological and physiological bases of *Ca*. Tisiphia-host interactions.

**Importance statement:** Members of the genus *Rickettsia* were originally identified as causative agents of mammalian vector-borne disease. In the last 25 years we have recognised that many *Rickettsia* are arthropod symbionts, and sit alongside a sister taxon, *Ca*. Megaira, which are symbiotic associates of microeukaryotes. The lack of genomic information for symbiotic strains affects our ability to determine the evolutionary relationships between strains and understand the biological underpinnings of the different symbioses. We clarify these relationships by assembling 26 genomes of *Rickettsia* from understudied groups, and the first two *Ca*. Megaira, from various insects and microeukaryotes. Of note, the accessory genome diversity and broad host range of Torix *Rickettsia* parallels all other *Rickettsia* combined. This diversity, alongside the breadth of host species, make the Torix clade an important hidden player in invertebrate biology and physiology. We argue this clade should be given its own genus status, for which we propose *Ca*. Tisiphia.

## Introduction

Symbiotic bacteria are vital to the function of most living eukaryotes, including microeukaryotes, fungi, plants, and animals (Boettcher et al., 1996; Clay et al., 2005; Douglas, 2011; Fujishima & Kodama, 2012). The symbioses formed are often functionally important to the host with effects ranging from mutualistic to detrimental. Mutualistic symbionts may provide benefits through the biosynthesis of metabolites, or by protecting their hosts against pathogens and parasitoids (Hendry et al., 2014; Oliver et al., 2010). Meanwhile parasitic symbionts can be detrimental to the host due to resource exploitation or through reproductive manipulation that favours its own transmission over the host’s (Engelstädter & Hurst, 2009; Leclair et al., 2017). Across these different symbiotic relationships, symbionts are often important determinants of host ecology and evolution.

The Rickettsiales (Alphaproteobacteria) represent an order of obligate intracellular bacteria that form symbioses with a variety of eukaryotes (Weinert et al., 2015). Within Rickettsiales, the family Rickettsiaceae represent a diverse collection of bacteria that infect a wide range of eukaryotic hosts and can act as symbionts, parasites, and pathogens. Perhaps the best-known clade of Rickettsiaceae is the genus *Rickettsia*, which was initially described as the cause of spotted fever and other rickettsioses in vertebrates that are transmitted by ticks, lice, fleas and mites (Angelakis & Raoult, 2017).

*Rickettsia* have been increasingly recognised as heritable arthropod symbionts. Since the first description of a maternally inherited male-killer in ladybirds (Werren et al., 1994), we now know that heritable *Rickettsia* are common in arthropods (Pilgrim et al., 2021; Weinert et al., 2009). Further, *Rickettsia*-host symbioses are diverse, with the symbiont capable of reproductive manipulation, nutritional and protective symbiosis, as well as influencing thermotolerance and pesticide susceptibility (Bodnar et al., 2018; Brumin et al., 2011; Chiel et al., 2009; Giorgini et al., 2010; Hurst et al., 1994; Kontsedalov et al., 2008; Łukasik et al., 2013).

Our understanding of the evolution and diversity of the genus *Rickettsia* and its allies has increased in recent years. Weinert et al. (2009) defined 13 different groups of *Rickettsia* with two early branching clades that appeared genetically distant from other members of the genus. The first of these was defined from a symbiont of *Hydra* and was named the Hydra group *Rickettsia*, which has since been assigned its own genus status, *Candidatus* Megaira (Schrallhammer et al., 2013). *Ca*. Megaira forms a sister clade to *Rickettsia* and is common in ciliate protists, amoebae, chlorophyte and streptophyte algae, and cnidarians (Lanzoni et al., 2019). Members of this clade are found in hosts from aquatic, marine and soil habitats which include model organisms (e.g., *Paramecium, Volvox*) and economically important vertebrate parasites (e.g., *Ichthyophthirius multifiliis*, the ciliate that causes white spot disease in fish) (Lanzoni et al., 2019). Whilst symbioses between *Ca*. Megaira and microeukaryotes are pervasive, there is no complete publicly available genome and the impact of these symbioses on the host are poorly understood.

A second early branching clade was first described from *Torix tagoi* leeches and is commonly coined Torix *Rickettsia* (Kikuchi & Fukatsu, 2005). Symbionts in the Torix clade have since been found in a wide range of invertebrate hosts from midges to freshwater snails, and in a fish-parasitic amoeba (Pilgrim et al., 2021). The documented diversity of hosts is wider than other *Rickettsia* groups, which are to date only found in arthropods and their associated vertebrate or plant hosts (Weinert et al., 2009). Torix clade *Rickettsia* are known to be heritable symbionts, but their impact on host biology is poorly understood, despite the economic and medical importance of several hosts (inc. bed bugs, black flies, and biting midges). Rare studies have described the potential effects on the host, which include: larger body size in leeches (Kikuchi & Fukatsu, 2005); a small negative effect on growth rate and reproduction in bed bugs (Thongprem et al., 2020); and an association with parthenogenesis in *Empoasca* Leafhoppers (Aguin-Pombo et al., 2021).

Current data seems to suggest an emerging macroevolutionary scenario where the members of *Rickettsia*-Megaira clade originated as symbionts of microeukaryotes, before diversifying to infect invertebrate symbionts. The Torix *Rickettsia* retained a broad range of hosts from microeukaryotes to arthropods. The remaining members of the genus *Rickettsia* evolved to be arthropod heritable symbionts and vector-borne pathogens. However, a lack of genomic and functional information for symbiotic clades limits our understanding of evolutionary transitions within *Rickettsia* and its sister groups. No *Ca*. Megaira genome sequences are currently publicly available and of the 165 *Rickettsia* genome assemblies available on the NCBI (as of 29/04/21), only two derive from the Torix clade and these are both draft genomes. In addition, dedicated heritable symbiont clades of *Rickettsia*, such as the Rhyzobius group, have no available genomic data, and there is a single representative for the Adalia clade. Despite the likelihood that heritable symbiosis with microeukaryotes and invertebrates was the ancestral state for this group of intracellular bacteria, available genomic resources are heavily skewed towards pathogens of vertebrates.

In this study we establish a richer base of genomic information for heritable symbiont *Rickettsia* and *Ca*. Megaira, then use these resources to clarify the evolution of these groups. We broaden available genomic data through a combination of targeted sequencing of strains without complete genomes, and metagenomic assembly of *Rickettsia* strains from arthropod genome projects. We report the first closed circular genome of a *Ca*. Megaira symbiont from a streptophyte alga (*Mesostigma viride*) and provide a draft genome for a second *Ca*. Megaira from a chlorophyte (*Carteria cerasiformis*). In addition, we present the first complete genomes of two Torix *Rickettsia* from a midge (*Culicoides impunctatus*) and a bed bug (*Cimex lectularius*) as well as a draft genome for *Rickettsia* from a tsetse fly (*Glossina morsitans submorsitans*, an important vector species), and a new strain from a spider mite (*Bryobia graminum*). A metagenomic approach established a further 22 draft genomes for insect symbiotic strains, including the first Rhyzobius and Meloidae group draft genomes. We utilize these to carry out pangenomic, phylogenomic and metabolic analyses of our extracted genome assemblies, with comparisons to existing *Rickettsia*.

## Methods

### Genomic data collection and construction

We employed two different workflows to assemble genomes for *Ca*. Megaira and *Rickettsia* symbionts (Figure 1). A) Targeted sequencing and assembly of focal *Ca*. Megaira and Torix *Rickettsia*. B) Assembly from SRA deposits of *Ca*. Megaira from *Mesostigma viride* NIES296 and the 29 arthropods identified in Pilgrim et al (2021) that potentially harbour *Rickettsia*. These were analysed alongside previously assembled genomes from the genus *Rickettsia*, and the outgroup taxon *Orientia tsutsugamushi*.

**Figure 1.**
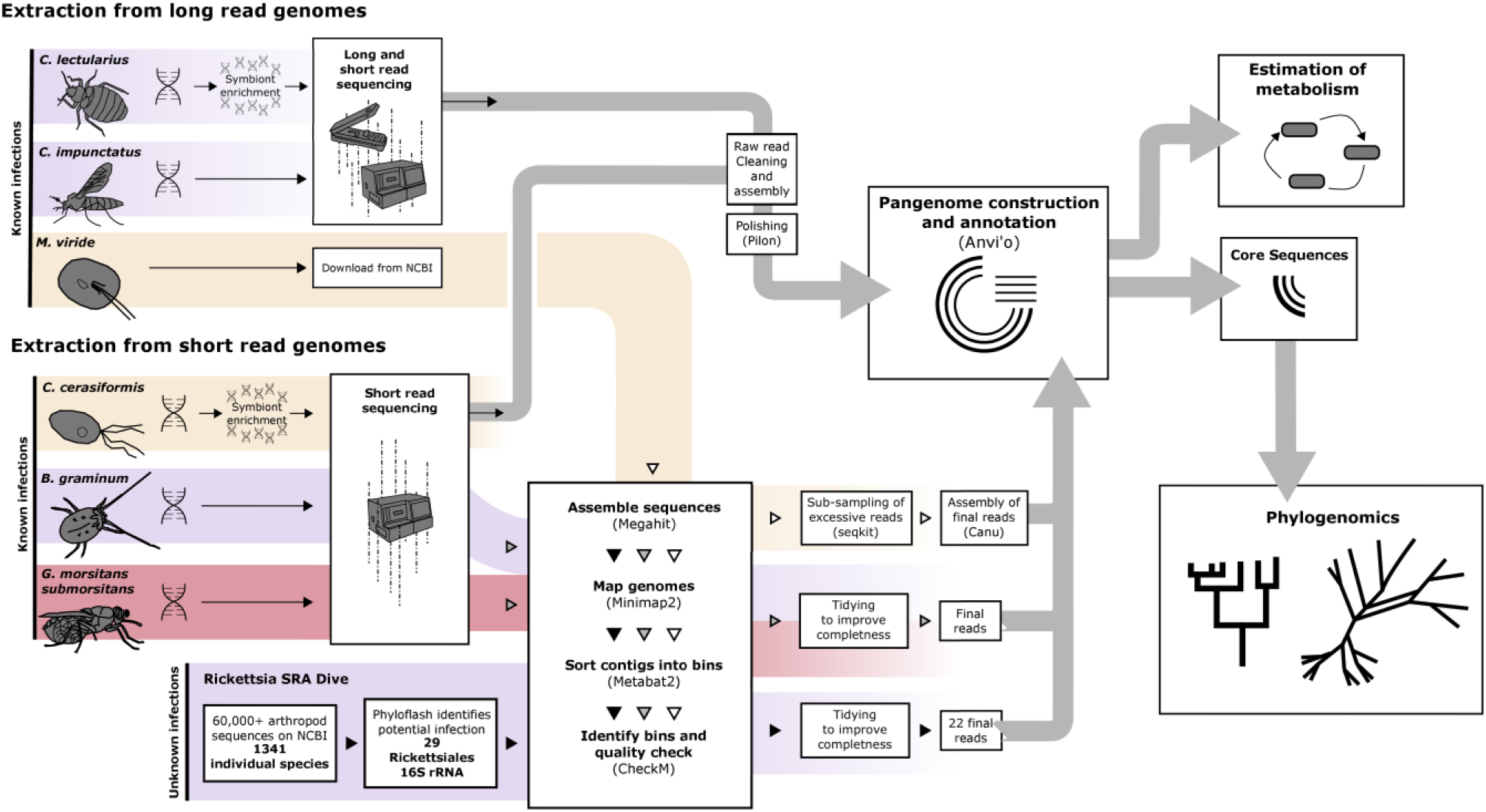
*Workflow diagram for extraction, assembly and analyses performed in this study. Purple highlights* Torix *Rickettsia and orange highlights Ca*. Megaira *and red highlights Transitional Rickettsia. A full resolution version can be found here: https://doi.org/10.6084/m9.figshare.15081975*.

DNA preparation, sequencing strategies and symbiont assembly methodologies varied between species. Methods are summarised in Figure 1 and detailed in supplementary material https://doi.org/10.6084/m9.figshare.14865582. The exact pipeline used to assemble genomes from Short Read Archive (SRA) data can be found here: https://figshare.com/s/d1155765b523a6379443.

#### Sample collection for targeted genome assembly

*Cimex lectularius* were acquired from the ‘S1’ isofemale colony maintained at the University of Bayreuth described in Thongprem et al (2020). *Culicoides impunctatus* females were collected from a wild population in Kinlochleven, Scotland (56° 42’ 50.7’’N 4° 57’ 34.9’’W) on the evenings of the 2nd and 3rd September 2020 by aspiration. *Carteria cerasiformis* strain NIES 425 was obtained from the Microbial Culture Collection at the National Institute for Environmental Studies, Japan. The *Glossinia morsitans submorsitans* specimen Gms8 was collected in Burkina Faso in 2010 and *Rickettsia* infection was present alongside other symbionts as described in Doudoumis et al. (2017). The assembly itself is a result of later thesis work (Blow, 2017).

A *Bryobia* mite community was sampled from herbaceous vegetation in Turku, Finland. The Moomin isofemale line was established by isolating a single adult female and was maintained on detached leaves of *Phaseolus vulgaris* L. cv Speedy at 25 °C, 60 % RH, and a 16:8 light:dark photoperiod. The Moomin spider mite line was morphologically identified as *Bryobia graminum* by Prof Eddie A. Ueckermann (North-West University).

#### Previously published Rickettsia genomes

A total of 86 published *Rickettsia* genomes, and one genome from *Orientia tsutsugamushi* were retrieved from the European Nucleotide Archive and assessed with CheckM v1.0.13 (Parks et al., 2015). Inclusion criteria for genomes were high completeness (CheckM > 90%), low contamination (CheckM < 2%) and low strain heterogeneity (Check M < 50%) except in the case of Adalia for which there is only one genome (87.6% completeness). Filtering identified 76 high quality *Rickettsia* genomes that were used in all subsequent analyses (S1 https://figshare.com/s/198c88c6e3ea5553192e).

### Genome content comparison and pangenome construction

Anvi’o 7 (Eren et al., 2021) was used to construct a pangenome for *Rickettsia*. Included in this were the 22 MAGs retrieved from SRA data, 2 *Ca*. Megaira genomes and 4 targeted Torix *Rickettsia* genomes, and one transitional group *Rickettsia* genome acquired in this study. To these were added the 76 published and 1 Orientia described above, giving a total of 104 genomes. Individual Anvi’o genome databases were additionally annotated with HMMER, KofamKOALA, and NCBI COG profiles (Aramaki et al., 2020; Eddy, 2011; Galperin et al., 2021). For the pangenome itself, orthologs were identified with NCBI blast, mcl inflation was set to 2, and minbit was 0.5. Genomes were arranged according to cluster presence absence and average nucleotide sequence identity was calculated using pyANI (Pritchard et al., 2016). See https://figshare.com/s/d1155765b523a6379443 for the exact code used in this section.

KofamKOALA annotation (Aramaki et al., 2020) in Anvi-o 7 was used to estimate completeness of metabolic pathways. Then Pheatmap (Kolde, 2019) in R 3.4.4 (R Core Team, 2020) was used to produce heatmaps of metabolic potential (figure 7). Annotations for function and *Rickettsia* group were added *post hoc* in Inkscape.

The biotin operon found in the genome Rhyzobius *Rickettsia* Oopac6 was identified from metabolic prediction (figure 7). To confirm Oopac6 carries a complete biotin pathway that shares ancestry with the existing *Rickettsia* biotin operon, Oopac6 biotin was compared to biotin pathways from five other related symbionts: *Cardinium, Lawsonia, Buchnera aphidicola, Rickettsia buchneri*, and *Wolbachia* (Seemann, 2014). Clinker (Gilchrist & Chooi, 2021) with default options was used to compare and visualise the similarity of genes within the biotin operon region of all 6 bacteria.

All generated draft and complete reference genomes were annotated using the NCBI’s Prokaryotic Genome Annotation Pipeline (PGAP) (Tatusova et al., 2016). Secondary metabolite biosynthetic gene clusters were identified using AntiSMASH version 6.0 (Blin et al., 2021) along with Norine (Flissi et al., 2019) which searched for similarities to predicted non-ribosomal peptides.

Functional enrichment analyses between the main *Rickettsia* clade and the Torix – *Ca*. Megaira clades were performed using the Anvi’o program anvi-get-enriched-functions-per-pan-group and the “COG_FUNCTION” as annotation source. A gene cluster presence - absence table was exported using the command “anvi-export-tables”. This was used to create an UpSet plot using the R package ComplexUpset (Krassowski et al., 2020; Lex et al., 2014) to visualize unique and shared gene clusters between different *Rickettsia* groups. A gene cluster was considered unique to a specified *Rickettsia* group when it was present in at least one genome belonging to that group. Gene cluster accumulation curves were performed for the pan-, core- and unique-genomes based on the same presence-absence matrix using a custom-made R script (Siozios, 2021). In each case the cumulative number of gene clusters were computed based on randomly sampled genomes using 100 permutations. The analysis was performed separately for each of the five major *Rickettsia* groups as well as the complete *Rickettsia* dataset. Curves were plotted using the ggplot2 R package (Wickham, 2016).

All information on extra genomes can be found at https://doi.org/10.6084/m9.figshare.14865582 and the code pipeline employed can be found at https://figshare.com/s/d1155765b523a6379443.

### Phylogeny, Network, and recombination

The single-copy core of all 104 genomes was identified in Anvi’o 7 and is made up of 74 single-copy gene (SCG) clusters. Protein alignments from SCG were extracted and concatenated using the command “anvi-get-sequences-for-gene-clusters”. Maximum likelihood phylogeny was constructed in IQ-TREE v2.1.2 (Nguyen et al., 2015). Additionally, 43 ribosomal proteins were identified through Anvi’o 7 to test phylogenomic relationships. These gene clusters were extracted from the pangenome and used for an independent phylogenetic analysis SUPPLEMENTARY FIG. The best model according to the Bayesian Information Criterion (BIC) was selected with Model Finder Plus (MFP) (Kalyaanamoorthy et al., 2017) as implemented in IQ-TREE; this was JTTDCMut+F+R6 for core gene clusters and JTTDCMut+F+R3 for ribosomal proteins. Both models were run with Ultrafast Bootstrapping (1000 UF bootstraps) (Hoang et al., 2018) with *Orientia tsutsugamushi* as the outgroup.

The taxonomic placement of Oopac6, Ppec13 and Dallo3 genomes within the Rhyzobius, Meloidae and Belli groups respectively were confirmed in a smaller phylogenetic analysis, performed as detailed in (Pilgrim et al. 2021) using reference MLST sequences (*gltA*, 16s rRNA, 17kDa, *COI*) from other previously identified *Rickettsia* profiles (S1 https://figshare.com/s/198c88c6e3ea5553192e). The selected models used in the concatenated partition scheme were as follows: 16S rRNA: TIM3e+I+G4; 17KDa: GTR+F+I+G4; COI: TPM3u+F+I+G4; gltA: K3Pu+F+I+G4a.

A nearest neighbour network was produced for core gene sets with default settings in Splitstree4 to further assess distances and relationships between *Rickettsia, Ca*. Megaira and Torix clades. All annotation was added post hoc in Inkscape. Furthermore, recombination signals were examined by applying the Pairwise Homoplasy Index (PHI) test to the DNA sequence of each core gene cluster extracted with Anvio-7. DNA sequences were aligned with MUSCLE (Edgar, 2004) and PHI scores calculated for each of the 74 core gene cluster with PhiPack (Bruen et al., 2006).

The taxonomic identity for new and newly expanded groups was established with GTDB-Tk (Chaumeil et al., 2020) to support the designation of new taxa through phylogenetic comparison of marker genes against an online reference database.

## Results and Discussion

We have expanded the available genomic data for several *Rickettsia* groups through a combination of draft and complete genome assembly. This includes an eight-fold increase in available Torix-group genomes, and the first available genomes for Meloidae and Rhyzobius groups. We further report the first reference genomes for *Ca*. Megaira.

### Complete and closed reference genomes for Torix *Rickettsia* and *Ca*. Megaira

The use of long-read sequencing technologies produced the first complete genomes for two subclades of the Torix group (RiCimp-limoniae, RiClec-leech). Sequencing depth of the *Rickettsia* genomes from *C. impunctatus* (RiCimp) and *C. lectularius* (RiClec) were 18X and 52X respectively. The RiCimp genome provides the first evidence of plasmids in the Torix group (pRiCimp001 and pRiCimp002). In addition, we assembled the first complete closed reference genome of *Ca*. Megaira from *Mestostigma viride* (MegNEIS296) from previously published genome sequencing efforts.

General features of both genomes are consistent with previous genomic studies of the Torix group (Table 1). A single full set of rRNAs (16S, 5S and 23S) and a GC content of ~33% was observed. Notably, the two complete Torix group genomes show a distinct lack of synteny (see S2 https://doi.org/10.6084/m9.figshare.14866263), a genomic feature that is compatible with our phylogenetic analyses that placed these two lineages in different subclades (leech/limoniae) (figures 2 and 3). Of note within the closed reference genomes MegNEIS296 and RiCimp, is the presence of a putative non-ribosomal peptide synthetase (NRPS) and a hybrid non-ribosomal peptide/polyketide synthetase (NRPS/PKS) respectively (see S3 https://doi.org/10.6084/m9.figshare.14865570). Although, the exact products of these putative pathways are uncertain, *in silico* prediction by Norine suggests close similarity with both cytotoxic and antimicrobial peptides hinting at a potential defensive role (see S3 https://doi.org/10.6084/m9.figshare.14865570). A hybrid NRPS/PKS cluster has previously been reported in *Rickettsia buchneri* on a mobile genetic element, providing potential routes for horizontal transmission (Hagen et al., 2018). In addition, putative toxin-antitoxin systems similar to the one associated with cytoplasmic incompatibility in *Wolbachia* have recently been observed on the plasmid of *Rickettsia felis* in a parthenogenetic booklouse (Gillespie et al., 2015, 2018). Toxin-endotoxin systems are thought to be part of an extensive bacterial mobilome network associated with reproductive parasitism (Gillespie et al., 2018). A BLAST search found a very similar protein in Oopac6 to the putative large pLbAR toxin found in *R. felis* (88% aa identity), and a more distantly related protein in the *C. impunctatus* plasmid (25% aa identity).

**Table 1.** *Summary of the complete Ca*. Megaira *and* Torix *Rickettsia genomes*

| Group | *Ca. Megaira* | *Torix Rickettsia* |  |
| --- | --- | --- | --- |
| Sub-group |  | Leech | Limoniae |
| Strain Name | MegNIES296 | RiCimp | RiClec |
| Symbiont genome accession | CP084576-CP084577 | CP084573-CP084575 | CP084572 |
| Host | Mesostigma viride NIES-296 | Culicoides impunctatus | Cimex lectularius |
| Raw reads accession | SRR8439255,<br>SRX5120346 | SRR16018514,<br>SRR16018513 | SRR16018512,<br>SRR16018511 |
| Total nucleotides | 1,532,409 | 1,566,468 | 1,611,726 |
| Chromosome size (bp) | 1,448,425 | 1,469,631 | 1, 611,726 |
| Plasmids | 1 (83,984 bp) | 2 (77550bp + 19287bp) | None |
| GC content (%) | 33.9 | 32.9 | 32.8 |
| Number of CDS | 1,359 | 1,397 | 1,544 |
| Avg. CDS length (bp) | 998 | 900 | 874 |
| Coding density (%) | 88.5 | 86 | 84 |
| rRNAs | 3 | 3 | 3 |
| tRNAs | 34 | 34 | 35 |

**Figure 2.**
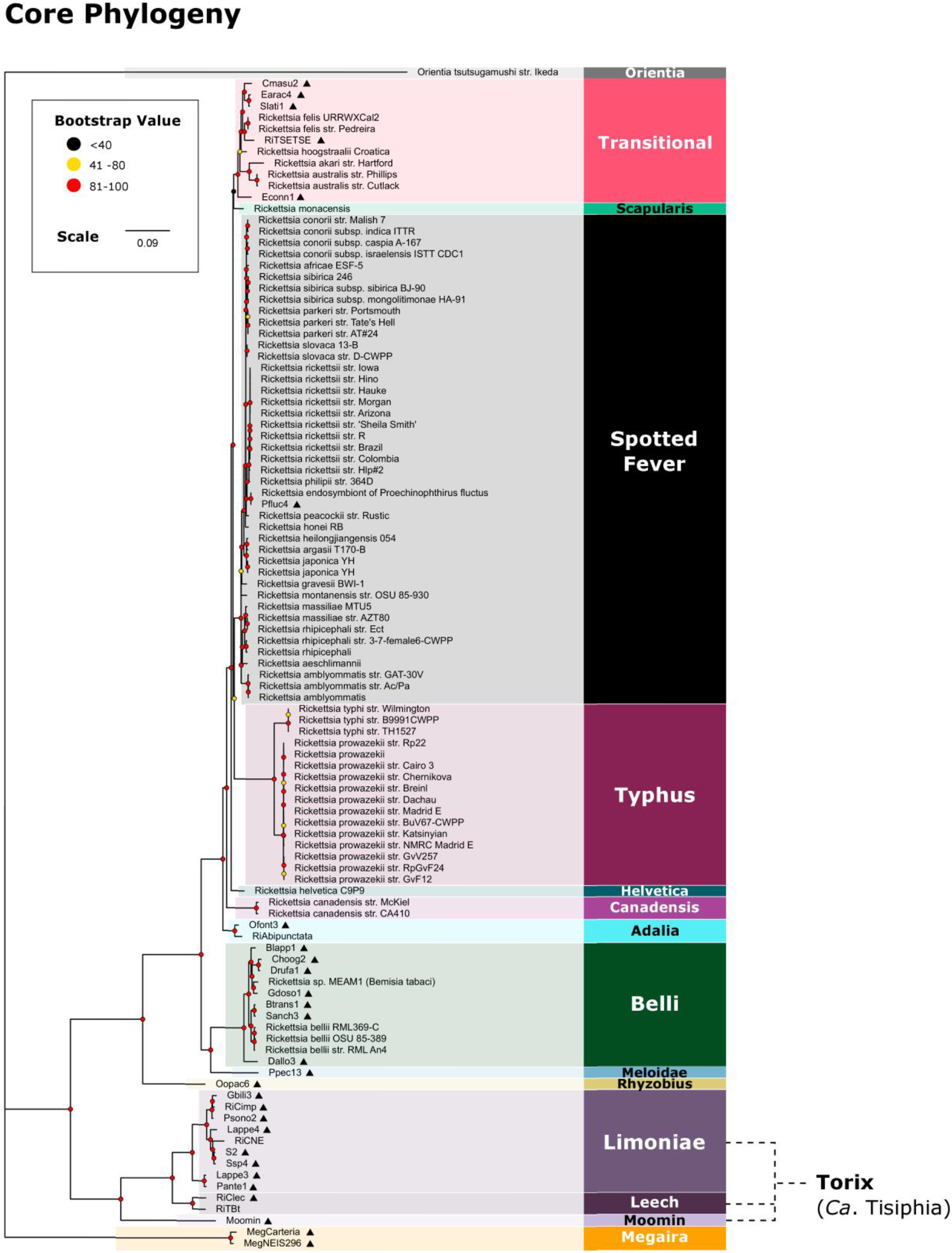
*Rickettsia and Ca*. Megaira *maximum likelihood (ML) phylogeny constructed from 74 core gene clusters extracted from the pangenome. New genomes are indicated by ▲ and bootstrap values based on 1000 replicates are indicated with coloured circles. New complete genomes are: RiCimp, RiClec and MegNEIS296. A full resolution version can be found here: https://doi.org/10.6084/m9.figshare.15081975*.

**Figure 3.**
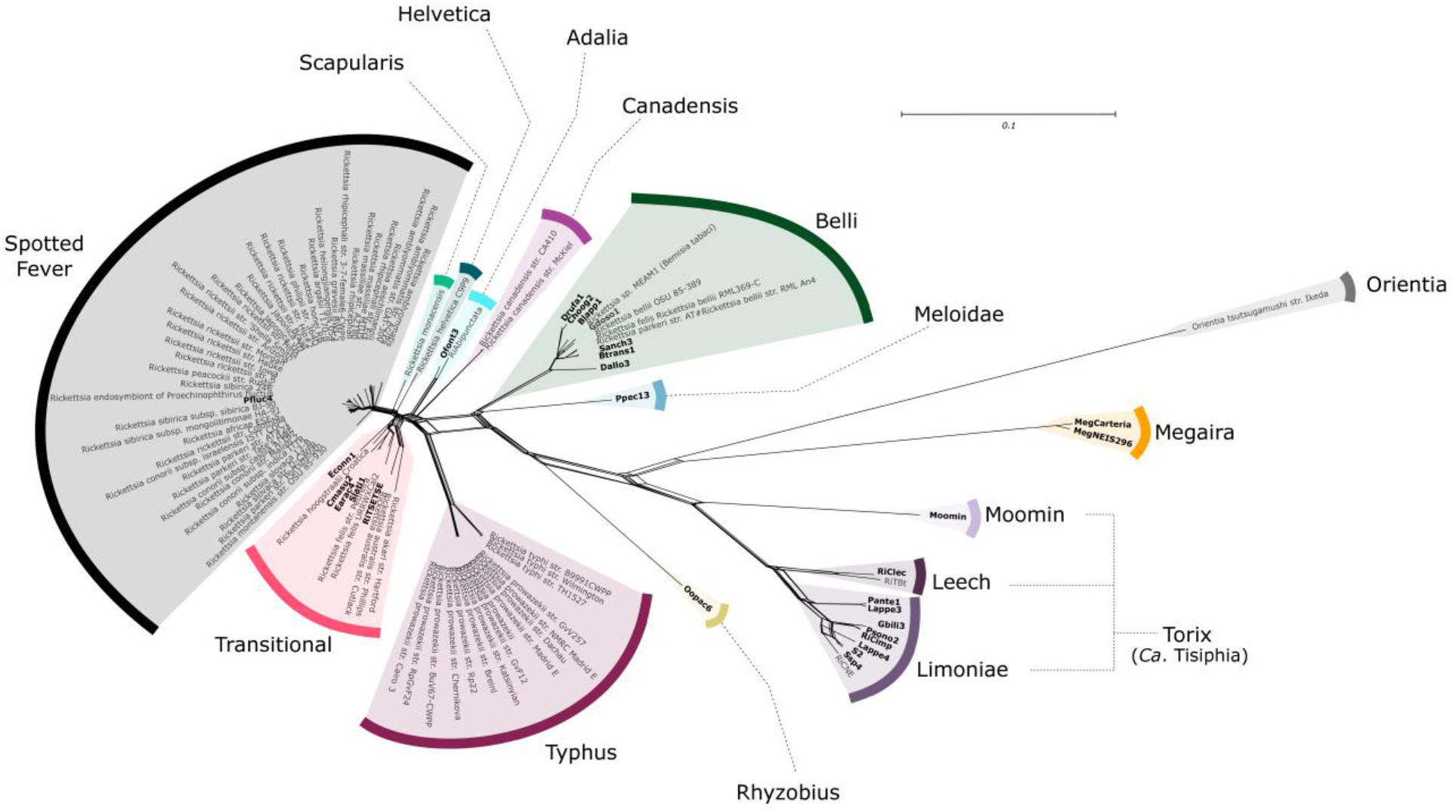
*Nearest Neighbour Network, displaying the distances between the 74 core gene sets across all 104 Rickettsia, Ca*. Megaira *genomes, and the outgroup Orientia. New genomes are indicated with bold text. A full resolution version can be found here: https://doi.org/10.6084/m9.figshare.15081975*.

**Figure 4.**
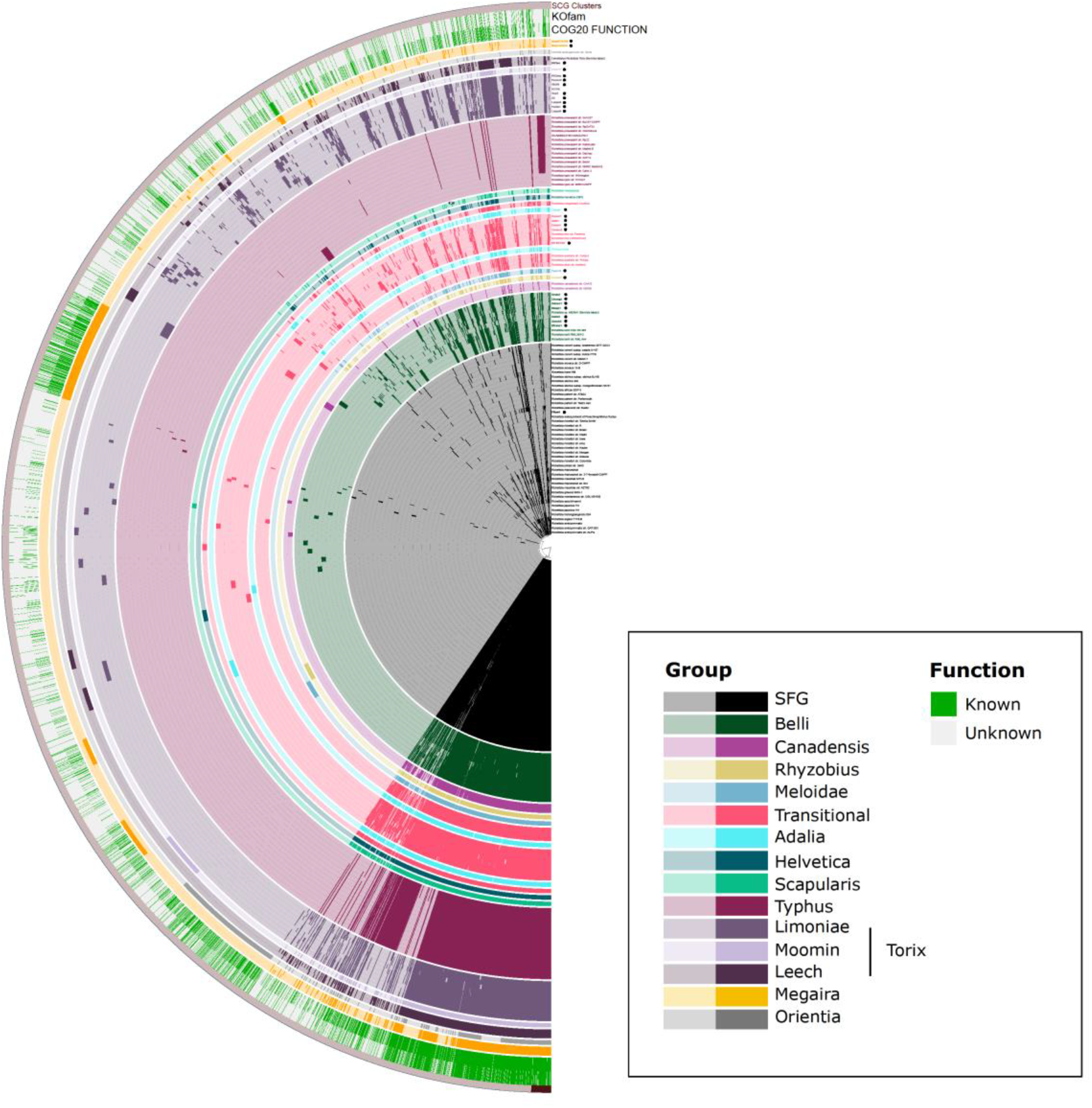
*Pangenome of all 104 genomes including Rickettsia, Torix, Ca*. Megaira *and the outgroup Orientia. New genomes are indicated by ●. Each genome displays gene cluster presence/absence and is organised by gene cluster frequency. Group identity was assigned from phylogeny. SFG is Spotted Fever Group. A full resolution version can be found here: https://doi.org/10.6084/m9.figshare.15081975*.

### Sequencing and *de novo* assembly of other *Rickettsia* and *Ca*. Megaira genomes

Our direct sequencing efforts enabled assembly of draft genomes for a second *Ca*. Megaira strain from the alga *Carteria cerasiformis*, and for *Rickettsia* associated with tsetse flies and *Bryobia* spider mites. The Transitional *Rickettsia* from a wild caught Tsetse fly, RiTSETSE, is a potentially chimeric assembly since we identified an excess of biallelic sites when the raw Illumina reads were mapped back to the assembly. It is also likely that RiTSETSE is not a heritable symbiont but comes from transient infection from a recent blood meal.

From the SRA accessions, the metagenomic pipeline extracted 29 full symbiont genomes for Rickettsiales across 24 host species. Five of 29 were identified as *Wolbachia* and discarded from further analysis, one was a *Rickettsia* discarded for low quality, and another was a previously assembled Torix *Rickettsia*, RiCNE (Pilgrim et al., 2017). Thus, 22 high quality *Rickettsia* metagenomes were obtained from 21 host species. One beetle (SRR6004191) carried coinfecting *Rickettsia* Lappe3 and Lappe4 (Table 2). The high-quality *Rickettsia* covered the Belli, Torix, Transitional, Rhyzobius, Meloidae and Spotted Fever Groups (Table 2 and S1 https://figshare.com/s/198c88c6e3ea5553192e).

**Table 2.** Brief summary of draft genomes generated during the current project and their associated hosts. Full metadata can be found in S1 https://figshare.com/s/198c88c6e3ea5553192e.

| Strain | Bacteria<br>Biosample<br>Accession | Group | Number<br>of<br>contigs | Total<br>length<br>(bp) | Host name | Order |
| --- | --- | --- | --- | --- | --- | --- |
| Blapp1 | SAMN21822536 | Belli | 171 | 1266633 | *Bembidion lapponicum* | Coleoptera |
| Btrans1 | SAMN21822537 | Belli | 241 | 1417452 | *Bembidion nr. transversale*<br>OSAC:DRMaddison<br>DNA3205 | Coleoptera |
| Choog2 | SAMN21822538 | Belli | 16 | 1357829 | *Columbicola hoogstraali* | Phthiraptera |
| Cmasu2 | SAMN21822539 | Transitional | 196 | 1295004 | *Ceroptres masudai* | Hymenoptera |
| Dallo3 | SAMN21822540 | Belli | 196 | 990679 | *Diachasma alloeum* | Hymenoptera |
| Drufa1 | SAMN21822541 | Belli | 14 | 1364611 | *Degeeriella rufa* | Phthiraptera |
| Earac4 | SAMN21822542 | Transitional | 96 | 1350066 | *Ecitomorpha arachnoides* | Coleoptera |
| Econn1 | SAMN21822543 | Transitional | 238 | 1070326 | *Eriopis connexa* | Coleoptera |
| Gbili3 | SAMN21822544 | Torix<br>limoniae | 171 | 1188102 | *Gnoriste bilineata* | Diptera |
| Gdoso1 | SAMN21822545 | Belli | 34 | 1420758 | *Graphium doson* | Lepidoptera |
| Lappe3 | SAMN21822558 | Torix<br>limoniae | 122 | 1368980 | *Labidopullus appendiculatus* | Coleoptera |
| Lappe4 | SAMN21822559 | Torix<br>limoniae | 154 | 1332357 | *Labidopullus appendiculatus* | Coleoptera |
| MegCarte-<br>ria | SAMN21822546 | Ca.<br>Megaira | 72 | 1298707 | *Carteria cerasiformis* | Chlamydomonadales |
| Ofont3 | SAMN21822560 | Adalia | 91 | 1529137 | *Omalius fontisbellaquei* | Coleoptera |
| Oopac6 | SAMN21822548 | Rhyzobius | 181 | 1497231 | *Oxypoda opaca* | Coleoptera |
| Pante1 | SAMN21822549 | Torix<br>limoniae | 70 | 1472610 | *Pseudomimeciton antennatum* | Coleoptera |
| Pfluc4 | SAMN21822550 | Spotted<br>Fever<br>Group | 7 | 1251895 | *Proechinophthirus fluctus* | Phthiraptera |
| Ppec13 | SAMN21822551 | Belli | 90 | 1426047 | *Pyrocoelia pectoralis* | Coleoptera |
| Psono2 | SAMN21822552 | Torix<br>limoniae | 163 | 1492063 | *Platyusa sonomae* | Coleoptera |
| RiTSETSE | SAMN21822553 | Transitional | 172 | 1451997 | *Glossina morsitans submorsitans* | Diptera |
| S2 | SAMN21822554 | Torix<br>limoniae | 103 | 1251484 | *Sericostoma* | Trichoptera |
| Sanch3 | SAMN21822555 | Belli | 181 | 1487154 | *Stiretrus anchorago* | Hemiptera |
| Slati1 | SAMN21822556 | Transitional | 109 | 1301763 | *Sceptobius lativentris* | Coleoptera |
| Ssp4 | SAMN21822557 | Torix<br>limoniae | 87 | 1231013 | *Sericostoma sp.* HW-2014 | Trichoptera |
| moomin | SAMN21822560 | Torix<br>moomin | 204 | 1137559 | *Bryobia graminum* | Trombidiformes |

Beetles, particularly rove beetle (Staphylinidae) species, appear in this study as a possible hotspot of *Rickettsia* infection. *Rickettsia* has historically been commonly associated with beetles, including ladybird beetles (*Adalia bipunctata*), diving beetles (*Deronectes* sp.) and bark beetles (*Scolytinae*) (Hurst et al., 1994; Küchler et al., 2009; Perlman et al., 2006; Weinert et al., 2009; Zchori-Fein et al., 2006). Though a plausible and likely hotspot, this observation needs be approached with caution as this could be an artefact of skewed sampling efforts.

All genome metadata and source information can be found here https://figshare.com/s/198c88c6e3ea5553192e.

### Phylogenomic analyses and taxonomic placement of newly assembled genomes

#### Phylogeny, network, and recombination

The network and phylogeny illustrate the distance of Torix from *Ca*. Megaira and other *Rickettsia*, along with an extremely high level of within-group diversity in Torix compared to any other group (Figures 2 and 3). The phylogenies generated using core genomes are consistent with previously identified *Rickettsia* and host associations using more limited genetic markers. For instance, Pfluc4 from *Proechinophthirus fluctus* lice is grouped on the same branch as a previously sequenced *Rickettsia* from a different individual of *P. fluctus*. Four of 22 genomes from the SRA screen are identified as Transitional, 1 is in Spotted Fever Group, 1 is Adalia, 8 are Belli and 7 are Torix limoniae. Targeted sequences were confirmed as: Torix limoniae (RiCimp), Torix leech (RiClec), Transitional (RiTSETSE), *Ca*. Megaira (MegCarteria and MegNEIS296), and a new Torix clade, Moomin (Moomin). The new Torix include one double infection giving a total of 10 new genomes across 9 potential host species. The double infection is found within the rove beetle *Labidopullus appendiculatus*, forming two distinct lineages, Lappe3 and Lappe4 (Fig 2 and 3).

In addition, the pre-existing *Rickettsia helvetica*, which is typically cited as a member of the Spotted Fever group as a result of its first description in 1993 (Beati et al., 1993; Chisu et al., 2017), seems to form its own group in all trees and networks (figure 2, 3 and https://doi.org/10.6084/m9.figshare.14865606). We conclude from this that *Rickettsia helvetica* is most similar to Scapularis group *Rickettsia*, but because it does not fall into the same clade in any tree or network, it is likely that the strain belongs to a distinct lineage of tick-borne *Rickettsia*.

We also report the first putative Rhyzobius *Rickettsia* genomes extracted from the staphylinid beetle *Oxypoda opaca* (Oopac6) and Meloidae *Rickettsia* from the firefly *Pyrocoelia pectoralis* (Ppec13). They have high completeness (S1 https://figshare.com/s/198c88c6e3ea5553192e), low pseudogenisation, and consistently group away from the other draft and completed genomes (Figures 2 and 3). MLST analyses demonstrate that these bacteria are most like the Rhyzobius and Meloidae groups described by Weinert *et al*. (2009) (see S5 https://doi.org/10.6084/m9.figshare.14865600). The pangenome and metabolic profile of this draft genome suggests that Meloidae is a sister group to Belli and that Rhyzobius *Rickettsia* is superficially similar to Belli and Transitional groups. The Rhyzobius-group symbiont is phylogenetically distant from most *Rickettsia* and is potentially a sister clade linking Torix and the main *Rickettsia* clades. Further genome construction will help clarify this taxon and its relationship to the rest of *Rickettsia*.

The sequencing data for the wasp, *Diachasma alloeum*, used here has previously been described to contain a pseudogenised nuclear insert of *Rickettsia* material, but not a complete *Rickettsia* genome (Tvedte et al., 2019). The construction of a full, non-pseudogenised genome with higher read depth than the insect contigs, low contamination (0.95%) and high completion (93.13%) suggests that these reads likely represent a viable *Rickettsia* infection in *D. alloeum*. However, these data do not exclude the presence of an additional nuclear insert. It is possible for a whole symbiont genome to be incorporated into the host’s DNA (Hotopp et al., 2007), and there are recorded partial inserts of *Ca*. Megaira genomes in the *Volvox carteri* genome (Kawafune et al., 2015). The presence of both the insert and symbiont need confirmation through appropriate microscopy methods.

Recombination is low within the core genomes of *Rickettsia* and *Ca*. Megaira, but may occur between closely related clades that are not investigated here. Across all genomes, the PHI score is significant in 6 of the 74 core gene clusters, suggesting putative recombination events. However, it is reasonable to assume that most of these may be a result of systematic error due to the divergent evolutionary processes at work across *Rickettsia* genomes. Patterns of recombination can occur by chance rather than driven by evolution which cannot be differentiated by current phylogenetic methods (Murray et al., 2016). The function of each respective cluster can be found at https://figshare.com/s/198c88c6e3ea5553192e.

### Gene content and pangenome analysis

#### Pangenome

Across all 104 genomes used in the pangenome analysis (figure 2, full data in S6 https://doi.org/10.6084/m9.figshare.14865576), Anvi’o identified 208 core gene clusters of which 74 are represented by single-copy genes. Bacterial strains of the different *Rickettsia* groups, especially the neglected symbiotic Rickettsiaceae, seem to have large, open pangenomes an indication of rapid evolution. As expected, the more genomes that are included in analyses, the smaller the core genome extracted.

Torix is a distinctly separate clade sharing less than 70% ANI similarity to any *Rickettsia* or *Ca*. Megaira genomes. It contains at least three groups that reflect its highly diverse niche in the environment (figure 5) (Jain et al., 2018; Pilgrim et al., 2021; Rodriguez-R et al., 2021). Torix has the most unique genes out of all the clades in this study followed by *Ca*. Megaira and Belli clades (figure 6). Rarefaction gene accumulation analysis suggest that Torix is the group where each additional genome included increases the pangenome repertoire to the greatest extent (figure 7). Torix group is thus more diverse in terms of genome content and size of the pangenome than other *Rickettsia* groups.

**Figure 5.**
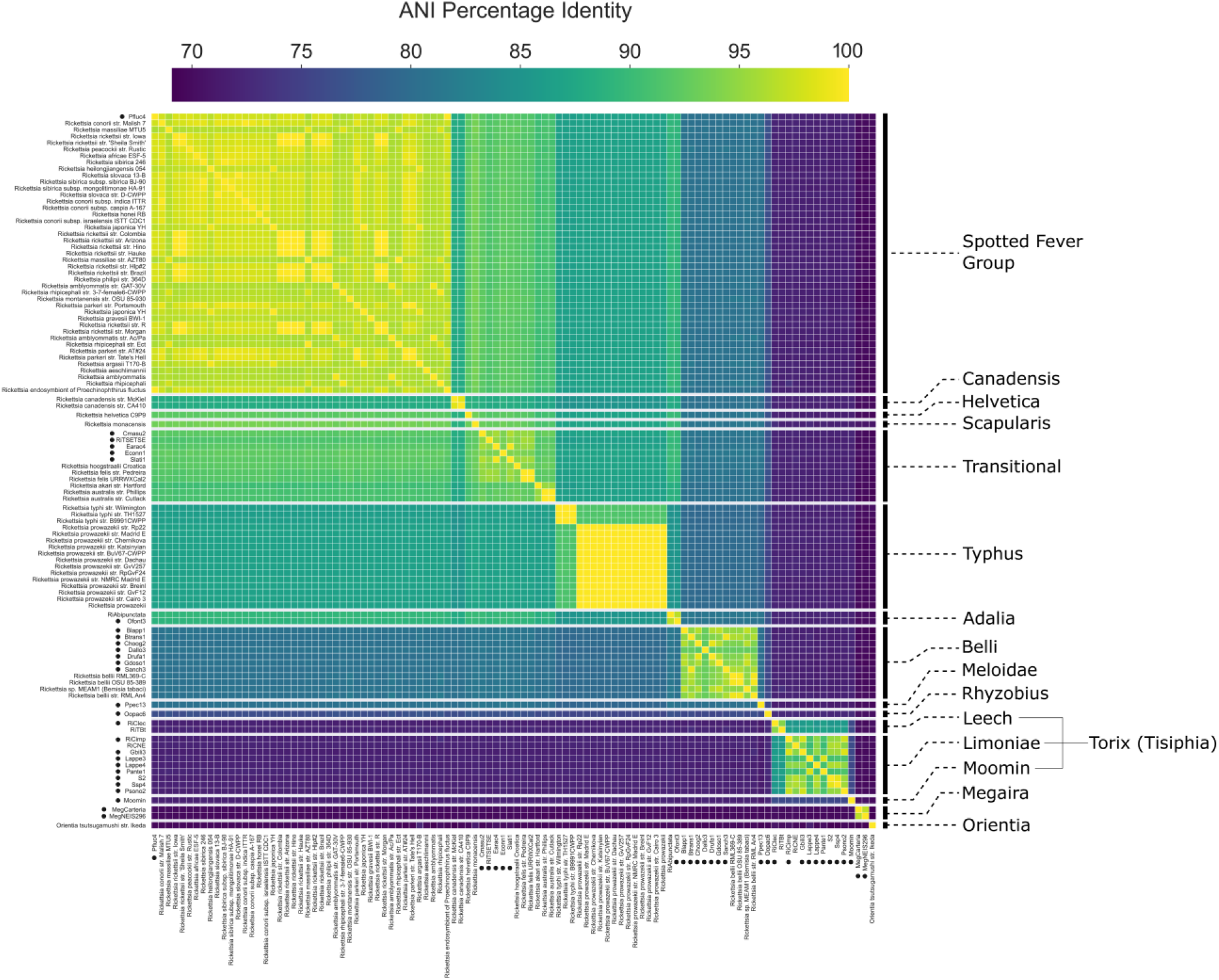
Pairwise Average Nucleotide Identity percentage across all genomes. New genomes are indicated by a black circle. A full resolution version can be found here: https://doi.org/10.6084/m9.figshare.15081975.

**Figure 6.**
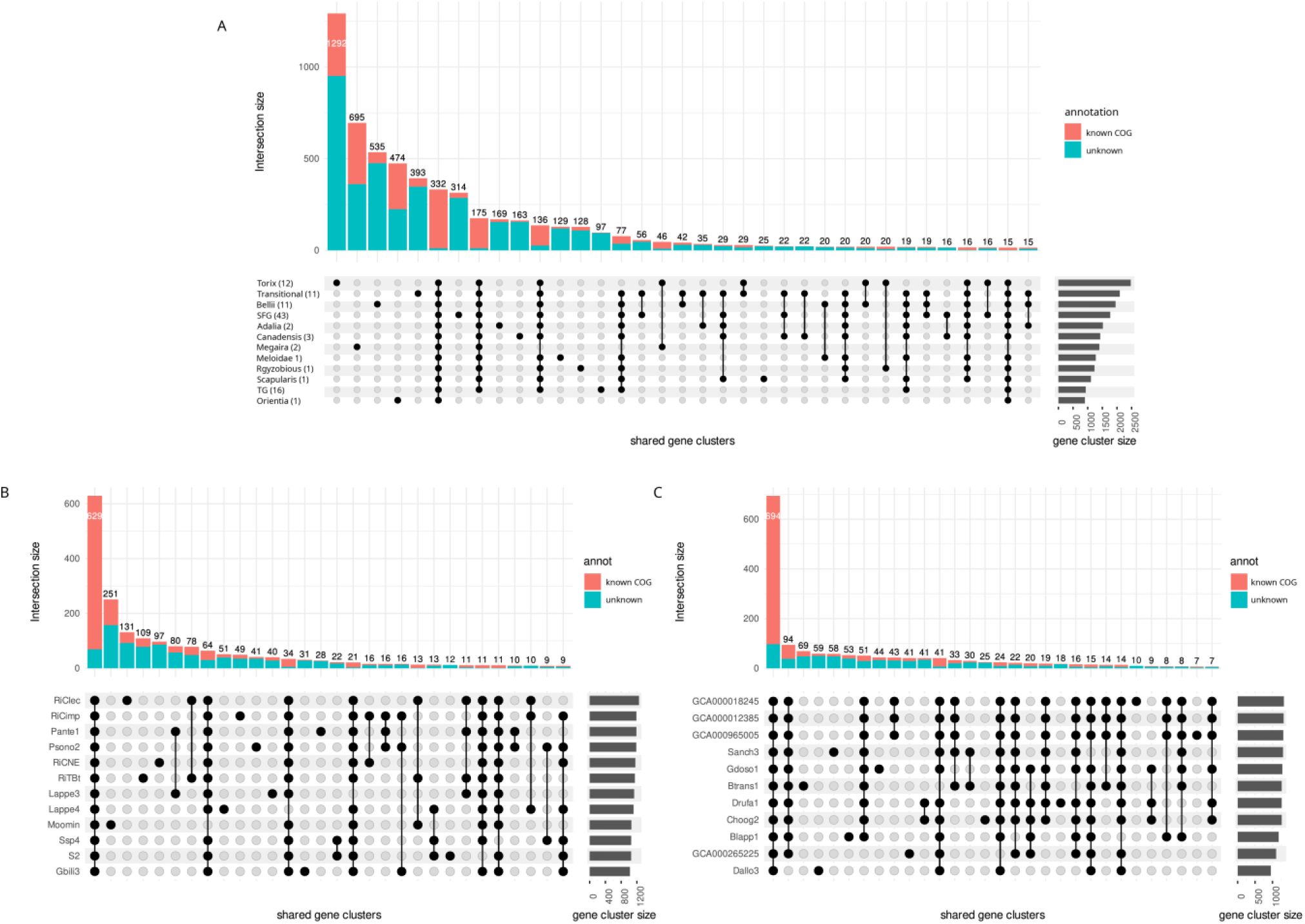
*Shared and unique gene clusters across A) All Rickettsia and Ca*. Megaira *genomes used in this study grouped by clade with Orientia as an outgroup B) all individual* Torix *genomes, and C) all individual Belli genomes. Horizontal grey bars to the right of each plot represent gene cluster size and vertical, coloured bars represent the size of intersections (the number of shared gene clusters) between genomes in descending order with known COG functions displayed in red and unknown in blue. Black dots mean the cluster is present and connected dots represent gene clusters that are present across groups. SFG is Spotted Fever Group and TG is Typhus Group. A full resolution version can be found here: https://doi.org/10.6084/m9.figshare.15081975*.

**Figure 7.**
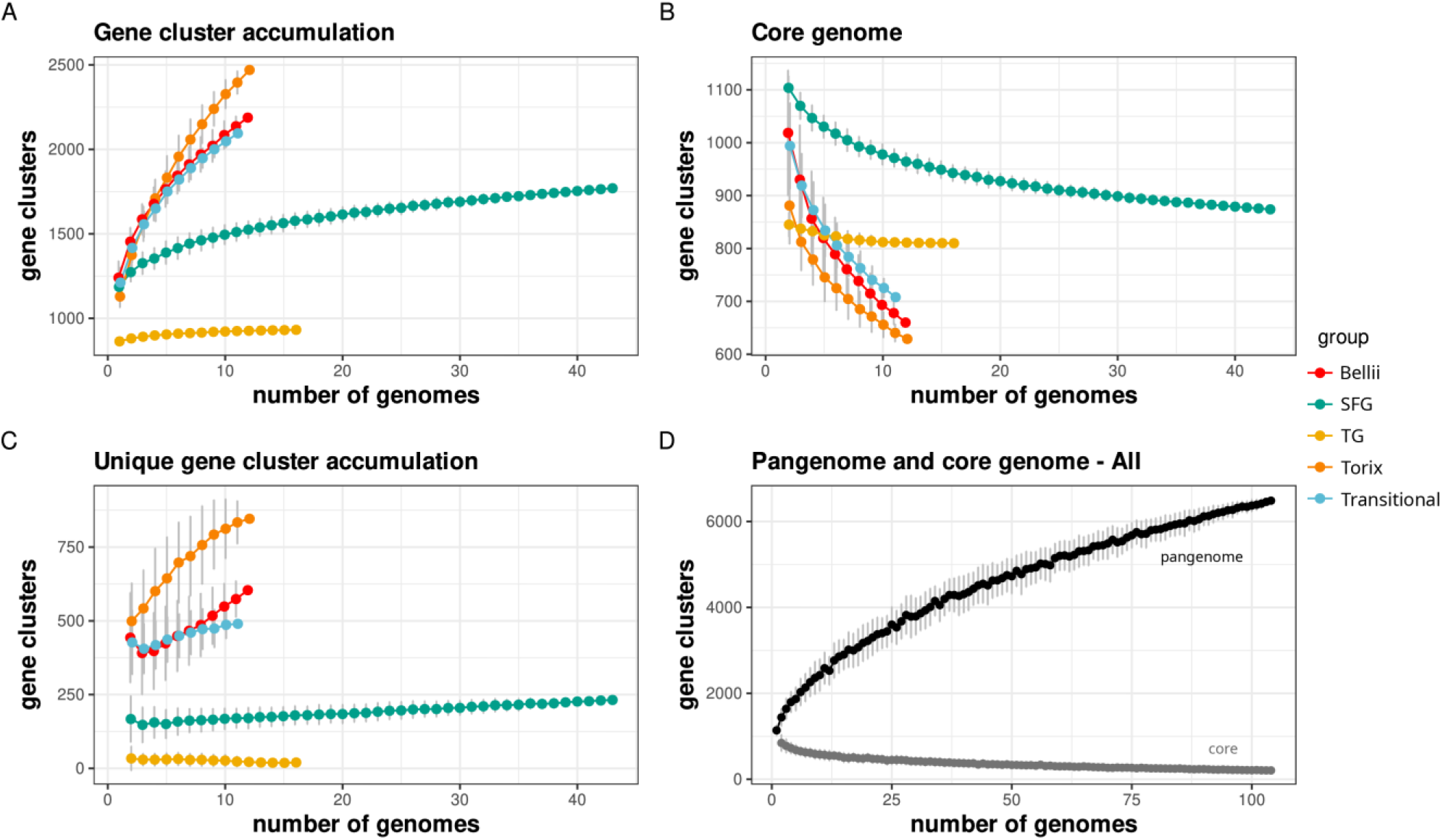
Gene cluster accumulation curves for pangenome (A), core genome (B) and the unique genome (C) of the 5 largest Rickettsia groups as a function of the number of genomes sequenced. The pangenome and the core genome accumulation curves for the complete Rickettsia dataset is shown in panel D. Error bars represent ± standard deviation based on 100 permutations. SFG is Spotted Fever Group and TG is Typhus Group. A full resolution version can be found here: https://doi.org/10.6084/m9.figshare.15081975.

*Rickettsia* lineages group together based on gene presence/absence and produce repeated patterns of accessory genes that reliably occur within each group (figure 2). ANI scores are also strongest within groups, while genomes tend to share lower similarity outside of their group (figure 4). This is particularly apparent in Torix and Ca. Megaira which are divergent from the main *Rickettsia* clade (figure 3 and 5).

#### Gene content and metabolic analyses

Rickettsial genomes extracted from SRA samples are generally congruent with the metabolic potential of their respective groups (Figure 8). Torix and *Ca*. Megaira have complete pentose phosphate pathways (PPP); a unique marker for these groups which seems to have been lost in the other *Rickettsia* clades. The PPP generates NADPH, precursors to amino acids, and is known to protect against oxidative injury in some bacteria (Christodoulou et al., 2018), as well as conversion of hexose monosaccharides into pentose used in nucleic acid and exopolysaccharide synthesis. The PPP has also been associated with establishing symbiosis between the Alphaproteobacteria *Sinorhizobium meliloti* and its plant host *Medicago sativa* (Hawkins et al., 2018). This pathway has previously been highlighted in Torix (Pilgrim et al., 2017) and its presence in all newly assembled Torix and *Ca*. Megaira draft genomes consolidates its importance as an identifying feature for these groups (Figure 8, S1 https://figshare.com/s/198c88c6e3ea5553192e). The PPP is likely an ancestral feature that was lost in the main *Rickettsia* clade.

**Figure 8.**
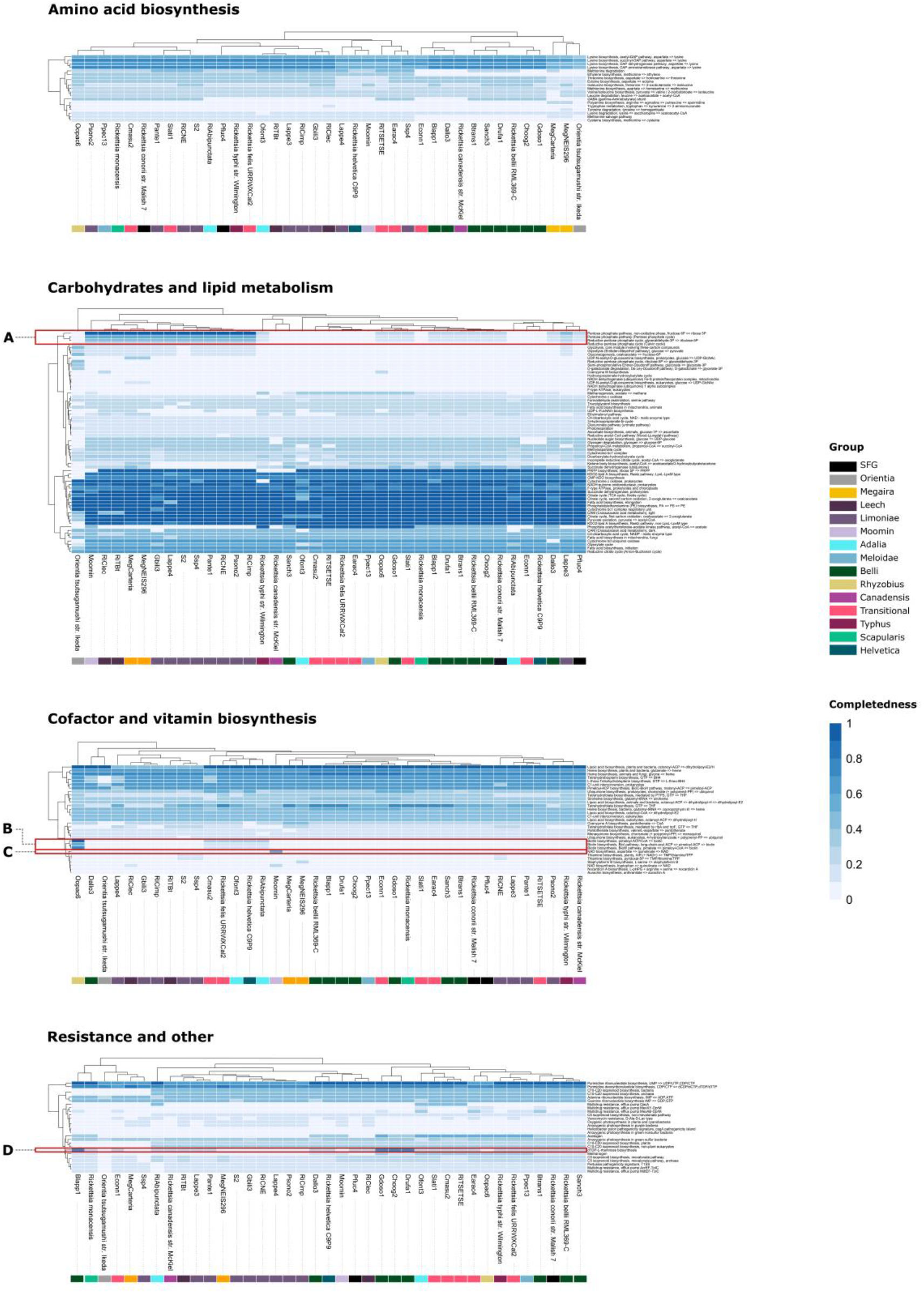
*Heatmaps of predicted KEGG pathway completion estimated in Anvi’o 7, separated by function and produced with Pheatmap. Pathways of interest are highlighted: A) The pentose phosphate pathway only present in* Torix *and Ca*. Megaira, *B) the biotin pathway present only in the Rhyzobius Rickettsia Oopac6, C) NAD biosynthesis only present in Moomin Rickettsia, D) dTDP-L-rhamnose biosynthesis pathway in Gdoso1, Choog2, Drufa1, and Blapp1. SFG is Spotted Fever. A full resolution version can be found here: https://doi.org/10.6084/m9.figshare.15081975*.

Glycolysis, gluconeogenesis and cofactor and vitamin metabolism are absent or incomplete across all *Rickettsia*, except the Rhyzobius group member, Oopac6 (Figure 8). Oopac6 has a complete biotin synthesis pathway that is related to, but distinct from, the *Rickettsia* biotin pathway first observed in *Rickettsia buchneri* (See S7 https://doi.org/10.6084/m9.figshare.14865567) (Gillespie et al., 2012). Based on the gene cluster comparison plot and an independent blastx search, the *GlyA* gene in *Rickettsia buchneri* appears to be a misidentified *bioF* gene (see S7 https://doi.org/10.6084/m9.figshare.14865567). Additionally, the insect SRA sample was not infected with *Wolbachia*, making it unlikely that the presence of the biotin operon is a result of misassembly. Animals can’t synthesize B-vitamins, so they either acquire them from diet or from microorganisms that can synthesize them. Oopac6 has retained or acquired a complete biotin operon where this operon is absent in other members of the genus. Biotin pathways in insect symbionts can be an indicator of nutritional symbioses (Douglas, 2017), so Rhyzobius *Rickettsia* could contribute to the feeding ecology of the beetle *O. opaca*. However, like other aleocharine rove beetles, *O. opaca* is likely predaceous, omnivorous or fungivorous (analysis of gut contents from a related species, *O. grandipennis*, revealed a high prevalence of yeasts: Klimaszewski et al., 2013). We posit no obvious reason for how these beetles benefit from harbouring a biotin-producing symbiont. One theory is that this operon could be a ‘hangover’ from a relatively recent host shift event and may have been functionally important in the original host. Similarly, if the symbiont is undergoing genome degradation, a once useful biotin pathway may be present but not functional (Blow et al., 2020). As this is the only member of this group with a complete genome so far, further research is required to firmly establish the presence and function of this pathway.

A 75% complete dTDP-L-rhamnose biosynthesis pathway was observed in 4 of the draft belli assemblies (Gdoso1, Choog2, Drufa1, Blapp1) (figure 8). Two host species are bird lice (*Columbicola hoogstraali, Degeeriella rufa*), one is a butterfly (*Graphium doson*), and one is a ground beetle (*Bembidion lapponicum*). dTDP-L-rhamnose is an essential component of human pathogenic bacteria like *Pseudomonas, Streptococcus* and *Enterococcus*, where it is used in cell wall construction (van der Beek et al., 2019). This pathway has also been utilized in the synthesis of plant cell walls (Jiang et al., 2021), may be involved in the moulting process of *Caenorhabditis elegans* (Feng et al., 2016), and is a precursor to rhamnolipids that are used in quorum sensing (Daniels et al., 2004). In the root symbiont *Azospirillium*, disruption of this pathway alters root colonisation, lipopolysaccharide structure and exopolysaccharide production (Jofré et al., 2004). No *Rickettsia* from typically pathogenic groups assessed in figure 8 has this pathway, and the hosts of these four bacteria are not involved with human or mammalian disease. Presence in feather lice provides little opportunity for this *Rickettsia* to be pathogenic as feather lice are chewers rather than blood feeders, and Belli group *Rickettsia* more generally are rarely pathogenic. Further, this association does not explain its presence in a butterfly and ground beetle; it is most likely that this pathway, if functional, would be involved in establishing infection in the insect host or host-symbiont recognition.

### Designation of *Ca*. Tisiphia

In all analyses, Torix consistently cluster away from the rest of *Rickettsia* as a sister taxon. Despite the relatively small number of Torix genomes, its within group diversity is greater than any divergence between previously described *Rickettsia* in any other group (figures 2, 3 and 5). Additionally, Torix shares characteristics with both *Ca*. Megaira and *Rickettsia*, but with many of its own unique features (figures 6 and 8). The distance of Torix from other *Rickettsia* and *Ca*. Megaira is confirmed in both the phylogenomic and metabolic function analyses to the extent that Torix should be separated from *Rickettsia* and assigned its own genus. This is supported by GTDB-Tk analysis which places all Torix genomes separate from *Rickettsia* (S1 https://figshare.com/s/198c88c6e3ea5553192e) alongside ANI percentage similarity scores less than 70% in all cases. To this end, we propose the name *Candidatus* Tisiphia after the fury Tisiphone, reflecting the genus *Ca*. Megaira being named after her sister Megaera.

## Conclusion

The bioinformatics approach has successfully extracted a substantial number of novel *Rickettsia* and *Ca*. Megaira genes from existing SRA data, including the first putative Rhyzobius *Rickettsia* and several *Ca. Tisiphia* (formerly Torix *Rickettsia*). Successful completion of two *Ca*. Megaira and two *Ca*. Tisiphia genomes provide solid reference points for the evolution of *Rickettsia* and its sister groups. From this, we can confirm the presence of a complete Pentose Phosphate Pathway in *Ca*. Tisiphia and *Ca*. Megaira, suggesting that this pathway was lost during *Rickettsia* evolution. We also describe the first Meloidae and Rhyzobius *Rickettsia* and show that Rhyzobius group *Rickettsia* has the potential to be a nutritional symbiont due to the presence of a complete biotin pathway. These new genomes provide a much-needed expansion of available data for symbiotic *Rickettsia* clades and clarification on the evolution of *Rickettsia* from *Ca*. Megaira and *Ca*. Tisiphia.

## Supporting information

All original genomes and raw readsets produced in this study can be accessed at Bioproject accession PRJNA763820 and all assemblies produced from previously published third party data can be accessed at Bioproject PRJNA767332.

Supplementary data and full resolution figures can be accessed on figshare here: https://doi.org/10.6084/m9.figshare.c.5518182.v1

## Acknowledgements

HRD was supported by the NERC ACCE Doctoral Training Programme. Grant code: NE/L002450/1 and NW was supported by a BOF post-doctoral fellowship (Ghent University, 01P03420) and by a Research Foundation - Flanders (FWO) Research Grant (1513719N). Funding for tsetse fly genomics were to ACD IP BBSRC projects BB/J017698/1 and BB/K501773/1, the materials from which were provided by P. Solano (Institut de Recherche pour le Développement, Unité Mixte de Recherche Interactions Hôtes-Vecteurs-Parasites-Environnement dans les Maladies Tropicales Négligées Dues aux Trypanosomatides, 34398, Montpellier, France) and J.-B. Rayaisse (Centre International de Recherche-Développement sur l’Élevage en zone Subhumide (CIRDES), N°559, Rue 5-31 Avenue du Gouverneur Louveau, 01 BP 454, Bobo Dioulasso 01, Burkina Faso). Jean-Baptiste died a few years ago but he was a fantastic person to work with and a great field entomologist. We also wish to thank Dr David Montagnes for teaching skills associated with algal culture.

We wish to thank Dr Débora Pires Paula (Embrapa) for granting permission to use SRA data for sample number SRR5651504, Iridian Genomes for allowing use of their SRA data, and the Microbial Culture Collection at the National Institute for Environmental Studies, Japan for use of the sample Carteria cerasiformis NIES-425.

## Contributions

Project concept: HRD, SS, JP and GH

Manuscript written by HRD, SS, JP and GDDH

SRA dive and metagenome assembly carried out by HRD with aid from SS.

Assembly of genome from SRA, pangenomics and phylogenomics carried out by HRD with advice from SS, GH

Metabolic analysis carried out by HRD, JP and SS

Sequencing and assembly of bacteria from Cimex lecticularius and Culicoides impunctatus genomes by SS and JP.

Sequencing and assembly of symbionts from Carteria by SHB and SS, supervised by PC and GH.

Sequencing and construction of RiTSETSE carried out by FB as part of thesis work supervised by AD.

SP collected and sequenced staphylinid genomes that were released through NCBI by iridian genomics.

NW collected and sequenced the *Bryobia* Moomin strain and performed preliminary metagenomic analyses

---

**S2.**
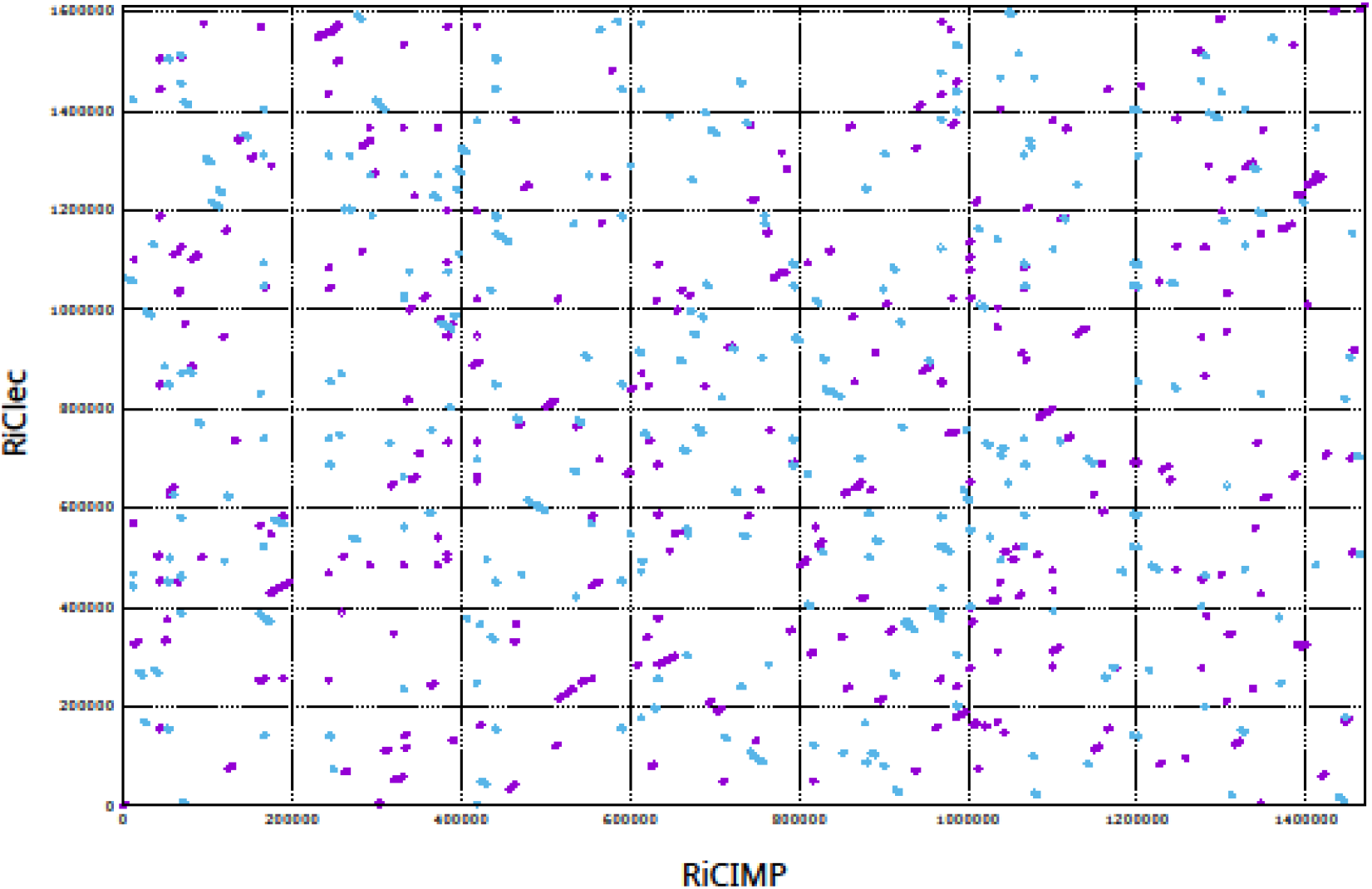
*Whole genome alignment between the complete* Torix *limoniae (RiCIMP) and* Torix *Leech (RiClec) genomes reveals complete lack of synteny. Magenta represents forward matches and blue reverse matches https://doi.org/10.6084/m9.figshare.14866263*.

**S3.**
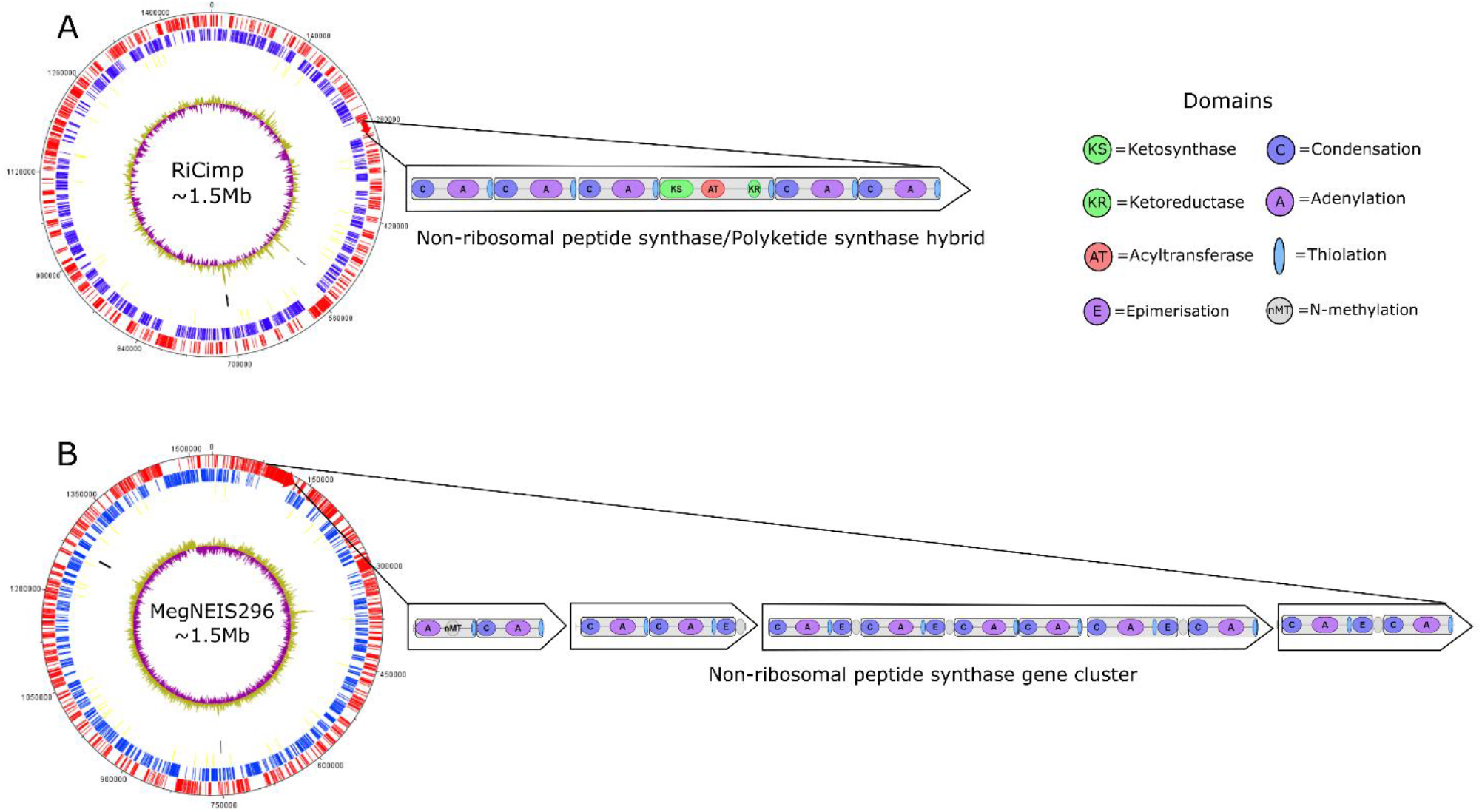
*The circular chromosomes of **A)** a* Torix *group Rickettsia (RiCimp) and **B)** a Ca*. Megaira *sp. (MegNEIS296). From outside to in, the circles represent: forward CDSs (Red), Reverse CDSs (blue), tRNAs (yellow) rRNAs (black), and GC content (green and magenta). Highlighted are the predicted modules formed by non-ribosomal peptide synthase genes (domains) that define individual amino acids in the synthesised peptide and show the catalytic domains within modules https://doi.org/10.6084/m9.figshare.14865570*.

**S4.**
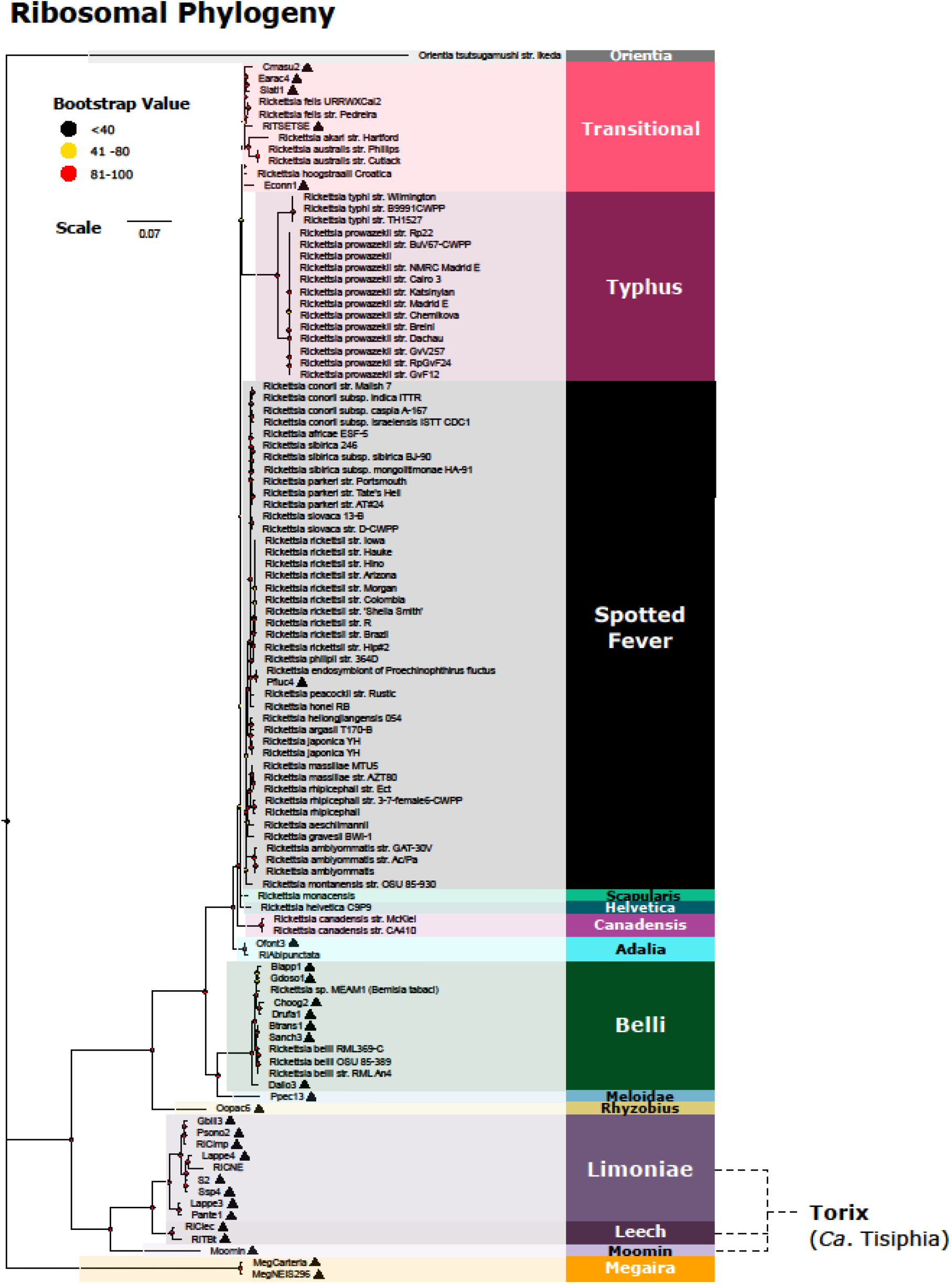
*Rickettsia and Ca*. Megaira *maximum likelihood (ML) phylogeny constructed from 43 ribosomal protein gene clusters extracted from the pangenome. New genomes are indicated by ▲ and bootstrap values based on 1000 replicates are indicated with coloured circles. New complete genomes are: RiCimp, RiClec and MegNEIS296. https://doi.org/10.6084/m9.figshare.14865606*

**S5.**
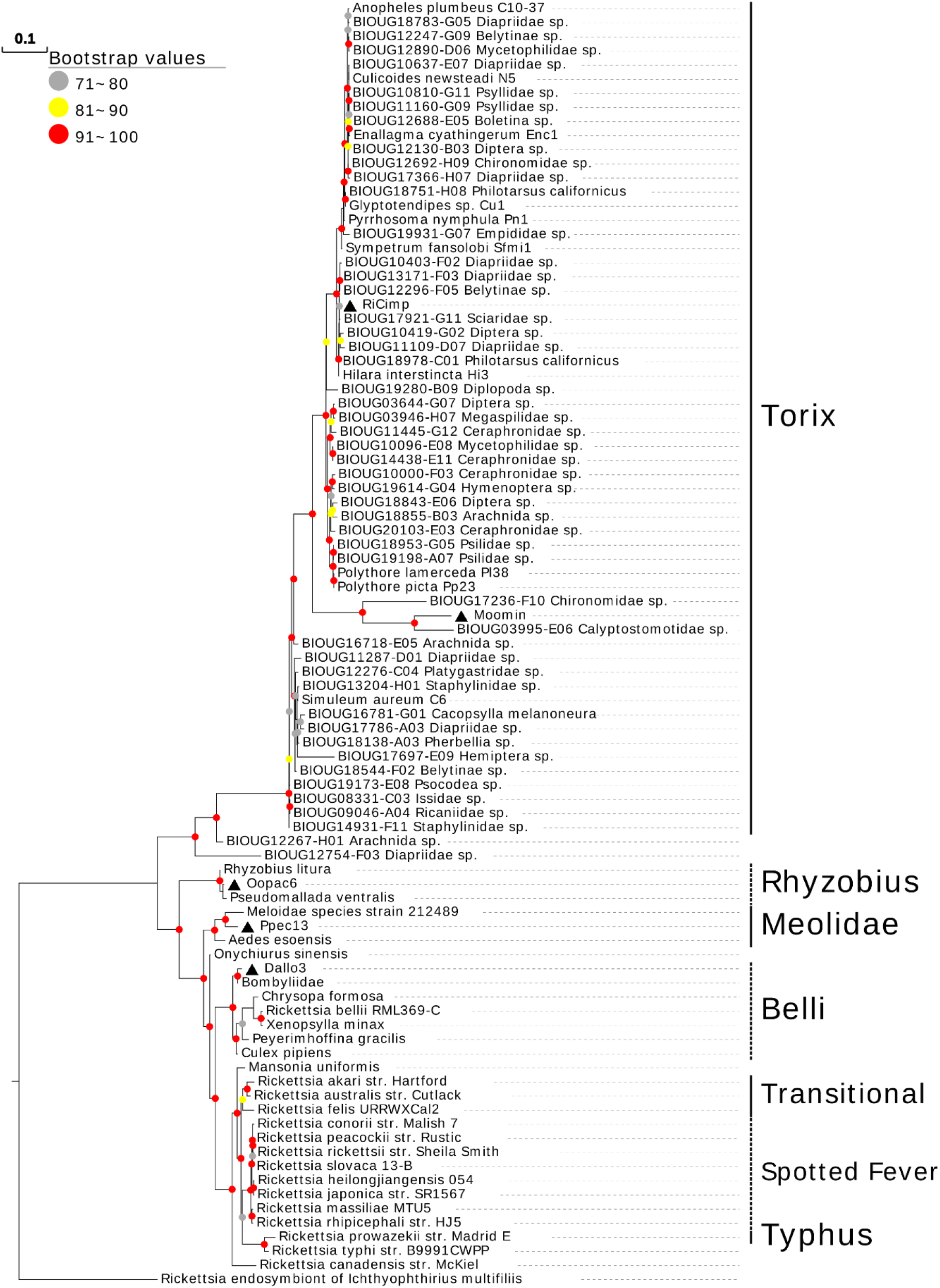
Phylogram of a maximum likelihood (ML) tree of 90 Rickettsia mutilocus profiles. The tree is based on 4 loci, 16S rRNA, 17KDa, gltA, and COI, under a partition model (2,781 bp total). https://doi.org/10.6084/m9.figshare.14865600

**S7.**
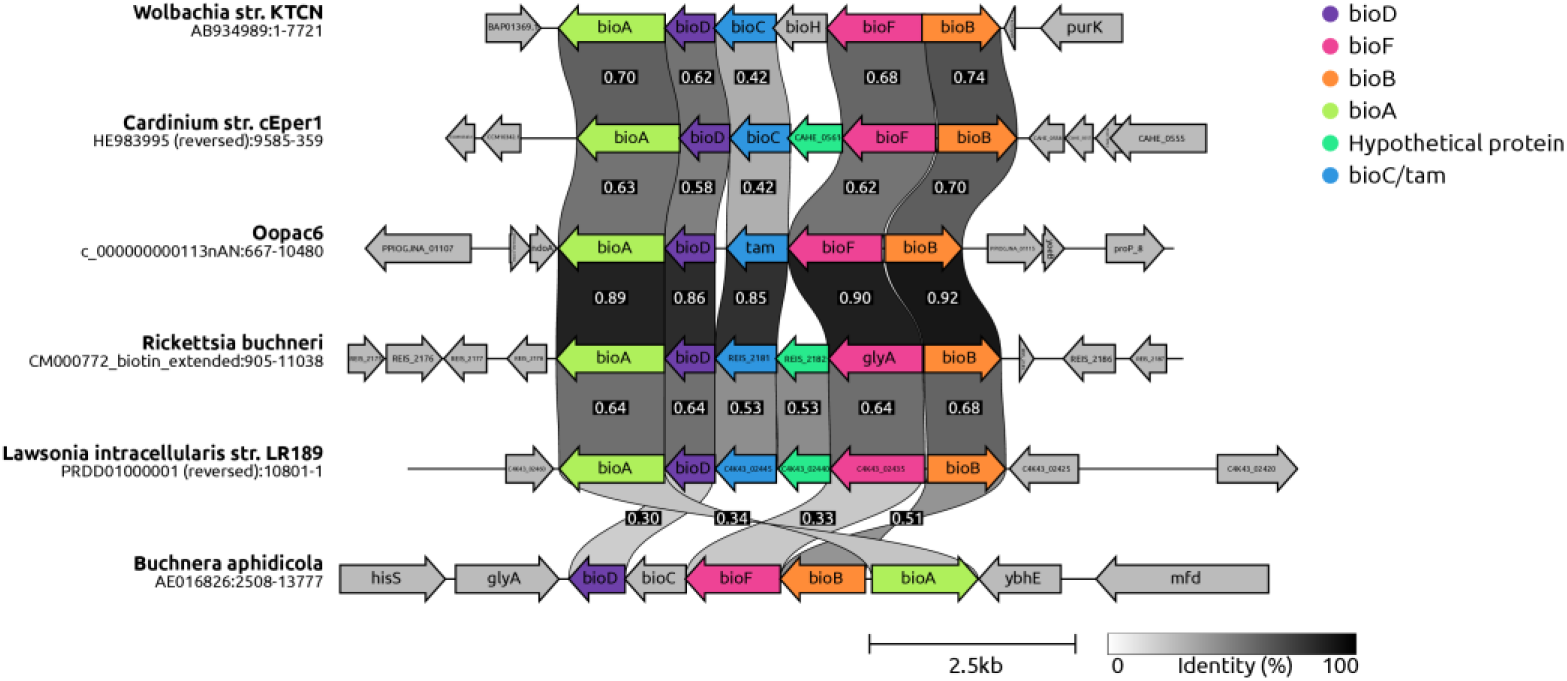
Biotin operon of the Rhyzobius Rickettsia, Oopac6, and its surrounding genes compared with other known biotin pathways in other related symbionts. Similarity scores in the black boxes refer to the percentage identity between the genes of the operon above or below it, further illustrated by a greyscale bar. Operons are ordered by overall similarity, showing the closest relationships between all 6. https://doi.org/10.6084/m9.figshare.14865567

